# Spatially and temporally distinct patterns of expression for VPS10P domain receptors in human cerebral organoids

**DOI:** 10.1101/2023.06.07.543827

**Authors:** Fabia Febbraro, Helena Hørdum Breum Andersen, Thomas E. Willnow

## Abstract

Vacuolar protein sorting 10 protein (VPS10P) domain receptors are a unique class of intracellular sorting receptors that emerge as major risk factors associated with psychiatric and neurodegenerative diseases, including bipolar disorders, autism, schizophrenia, as well as Alzheimer’s disease and frontotemporal dementia. Yet, the lack of suitable experimental models to study receptor functions in the human brain has hampered elucidation of receptor actions in brain disease. Here, we have adapted protocols in human cerebral organoids to the detailed characterization of VPS10P domain receptor expression during development and in mature brain tissue, including single-cell RNA sequencing. Our studies uncovered spatial and temporal patterns of expression unique to individual receptor species in the human brain. While *SORL1* expression is abundant in stem cells and *SORCS1* peaks at onset of neurogenesis, *SORT1* and *SORCS2* show increasing expression with maturation of neuronal and non-neuronal cell types, arguing for distinct functions in development versus the adult brain. In neurons, subcellular localization also distinguishes between two types of receptors, either mainly localized to the cell soma (*SORL1* and *SORT1*) or also to neuronal projections (*SORCS1* and *SORCS2*), suggesting divergent functions in protein sorting between Golgi and the endo-lysosomal system or along axonal and dendritic tracks. Taken together, our findings provide an important resource concerning temporal, spatial, and subcellular patterns of VPS10P domain receptor expression in cerebral organoids for further elucidation of receptor (dys)functions causative of behavioral and cognitive defects of the human brain.

## INTRODUCTION

Vacuolar protein sorting 10 protein (VPS10P) domain receptors are a specialized class of type-1 transmembrane receptors that sort target proteins between cell surface and intracellular compartments, defining endocytic and secretory capacities of cells. Mammalian members of the gene family encompass the sorting protein-related receptor with A-type repeats (SORLA), sortilin, as well as sortilin-related receptors CNS expressed (SORCS)-1, -2, and -3 (Fig. 1A) (reviewed in (Hermey, 2009; Malik and Willnow, 2020)). In line with prominent expression of VPS10P domain receptors in the central and peripheral nervous systems, these receptors have been implicated in multiple psychiatric and neurodegenerative disorders. Among others, SORCS1, -2 and -3 have been identified as major risk genes for bipolar disorder, attention deficit hyperactivity disorder (ADHD), autism, and schizophrenia (Baum et al., 2008; Christoforou et al., 2011; Lionel et al., 2011; Alemany et al., 2015). SORLA and sortilin are genetically associated with age-related neurodegeneration in Alzheimer’s disease (AD) (Rogaeva et al., 2007; Lambert et al., 2013; Bellenguez et al., 2022) and frontotemporal dementia (FTD) (Carrasquillo et al., 2010). The importance of VPS10P domain receptors as therapeutic targets for mental disorders is underscored by clinical trials using sortilin antagonists for the treatment of FTD (https://www.alzforum.org/therapeutics/latozinemab).

**Figure 1:**
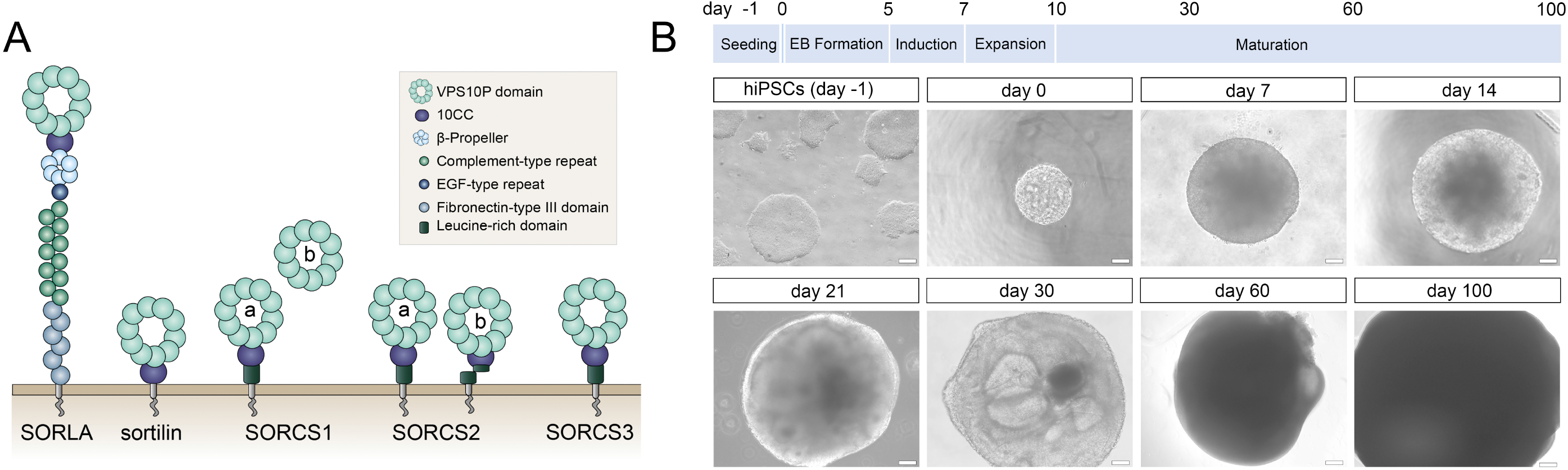
Human cerebral organoids as model system to study VPS10P domain receptors. (**A**) Structural organization of the mammalian VPS10P domain receptors sorting protein-related receptor with A-type repeats (SORLA), sortilin, as well as sortilin-related receptors CNS expressed (SORCS)-1, -2, and -3. These type-1 transmembrane proteins share a common vacuolar protein sorting 10 protein (VPS10P) domain that constitutes the main ligand binding site. Their ectodomains may harbor additional protein modules involved in protein-protein interactions. Their cytoplasmic tails contain binding motifs for sorting adaptors directing trafficking of the receptors between cell surface and intracellular compartments. For reasons of simplicity, all receptors are depicted as monomers. However, active forms of SORCS1, -2, -3 likely constitute homodimers, whereas sortilin transiently dimerizes in endosomes to discharge its ligands. Active SORCS1 exists as membrane-bound full-length receptor (version a) and as secreted ectodomain (version b). SORCS2 is found as single- (version a) and as proteolytically cleaved two-chain variants (version b) that each carry distinct receptor functions. (**B**) The timeline for differentiation of human induced pluripotent stem cells (hiPSCs) into cerebral organoids is given. The appearance of exemplary organoids at the indicated timepoints is shown in brightfield images below. EB, embryoid body. Scale bars: 100 µm or 200 µm (for day 60 and 100).

Much of our knowledge about the role of VPS10P domain receptors in brain health and disease stems from studies in mouse models with targeted disruption or induced overexpression of receptor genes (Malik and Willnow, 2020; Salasova et al., 2022). While these studies have been instrumental in identifying fundamental processes controlled by these receptors, they also have highlighted important differences in receptor biology in humans versus rodent models. For example, sortilin has been shown to promote lipoprotein particle secretion from mouse (Kjolby et al., 2010; Strong et al., 2012) but to block release from human cells (Musunuru et al., 2010). The role of SORLA as risk factor for both familial and sporadic forms of AD may be related to its expression in human microglia (Nott et al., 2019), yet relatively little expression in this cell type is seen in mice (Gosselin et al., 2017; Hansen et al., 2018). Obviously, validating the significance of single nucleotide polymorphisms (SNPs) associating VPS10P domain receptor genes with brain diseases also requires the use of human cell models derived from carriers of such risk gene variants.

Towards validating human iPSC-derived models for the study of VPS10P domain receptors in development and function of the human brain, we carried out a detailed analysis of the spatial and temporal aspects of receptor expression in human cerebral organoids.

## MATERIALS AND METHODS

### Cell lines

The induced pluripotent stem cell (iPSC) line HMGUi001 was used in this study (https://www.ebi.ac.uk/biosamples/samples/SAMEA5696688). It was reprogrammed from fibroblasts obtained from a Caucasian female donor. The cells were grown in Essential 8 Flex Basal Media (Gibco, A28583-01, Thermofisher) on vitronectin-coated plates (07180 Stemcell technologies; NUNC DISH 35 mm, 150255, Thermoscientific). The cells were passaged every 3 to 4 days at a ratio of 1:10 using 0.5 mM EDTA in PBS for 4 min at room temperature.

### Generation of cortical organoids

For generation of cerebral organoids, we adapted published protocols (Lancaster et al., 2013; Giandomenico et al., 2021). On day -1, iPSCs were resuspended in StemFlex media containing 10 µM Rock inhibitor Y-27632 (Cayman Chemical Company, Bionordika) and 2000 cells in 150 µl of media were plated in each well of 96-well ultra-low attachment plates (174929, Thermofisher). Cells were centrifuged at 100g for 3 min and incubated at 37°C (in 5% CO_2_) for 24 h. On day 0, 70 µl of media were removed from each well and replaced by 100 µl embryoid body (EB) formation medium (Cerebral organoid kit, cat. n. 08570, StemCell Technologies). On day 2, again 100 µl media were removed and the same amount of fresh EB formation media added back to each well. On day 5, 100 µl of media containing an organoid were removed from each organoid-containing 96-well using a pipette with 200 µl pipette tip. Once the residing organoids descended to the opening of the pipette tip by gravity, they were collected and moved to a new 96-well plate containing 150 µl of induction media (Cerebral organoid kit, cat. n. 08570, StemCell Technologies) per well. Special care was taken to only release the organoids from the tip by gravity and to exclude the contaminating EB medium. On day 7, 70 µl of media were removed from each well and replaced with 150 µl expansion medium (Cerebral organoid kit, cat. n. 08570, StemCell Technologies). On day 10, organoids were embedded in matrigel (Matrigel 356234 for organoids, BD Bioscience) as follows. Matrigel was thawed and kept on ice for at least 2 h before application to prevent polymerization. Using a 1 ml pipette tip, 32 organoids were collected from 96-well plates and placed in a 1.5 ml reaction tube. Once the organoids sunk to the bottom of the tube, the supernatant was removed and 60 µl of maturation medium were added, followed by 100 µl of Matrigel. The solutions were mixed gently, being careful not to stir up the organoids. Using a 200 µl pipette tip, the medium containing the organoids was recovered and plated in drops in ultra-low attachment 6-well plates (83.3920500, Starstedt). The plate was incubated at 37°C for 17 min, after which 4 ml maturation medium (Cerebral organoid kit, cat. n. 08570, StemCell Technologies) were gently added to each well. After 24 hours incubation, EB were detached from the bottom of the wells using a 1 ml pipette tip and the 6-well plate placed back into the incubator on an orbital shaker set to 74 rpm. Penicillin-Streptomycin solution (0.5%, Cat No. 15140122, Life and Technologies) was added for cell culture and differentiation. The medium was changed every other day and organoids were allowed to grow for up to 100 days. Samples were collected at different time points for further analysis.

### Single-cell sample preparation

Five to six 100 days old organoids were collected and placed in a 35 mm plate. To remove dead cells from the organoid core, organoids were cut in half using a syringe needle and washed with PBS. Subsequently, PBS was removed and 200 µl of TrypLE Express solution (12604-013 Gibco) were added. Next, the organoids were cut into smaller fragments using two needles attached to 1 ml syringes. Fragments were collected in a 1.5 ml reaction tube and subjected to gentle pipetting followed by intermittent resting at 37°C for 5 minutes. Single cells present in the supernatant after 5 min were collected into a new reaction tube and placed into the incubator. Tissue fragments remaining in the tube were treated with 100 µl of TrypLe Express and dispersed into small pieces using consecutive pipetting with 1 ml and 200 µl pipette tips. After additional five minutes, the single cells in the supernatant were again collected and added to the tube containing the single-cell suspension. This procedure was repeated 3 times for a total of 15 min. Any remaining tissue fragments were discarded. TrypLe Express dissociation was stopped by adding an equal volume of E8 Stem Flex medium, resulting in a total volume of 1 ml. Cell viability was assessed using the viability dye Trypan blue (0.4%, T10282, Invitrogen) and cells were counted using an automated cell counter (Countess II, Invitrogen). Thereafter, the single-cell suspension was filtered through a 40 µm filter (22363547, Thermo fisher) placed on top of a 50 ml tube. Filtered cells were collected and cell viability determined as above. Typically, cell viability at this step was around 50%-60%. To increase the percent of live cells in the single-cell suspension, the cells were centrifuged for 2 min at 200g and placed on ice. The cell pellet was washed in 1 ml of PBS with 0.04% BSA (130-091-376, Miltenyibiotec), pipetted gently to remove any cell clumps, and counted again. This washing step was repeated two more times, resulting in a cell viability of approximately 80%. Finally, 5000 cells were collected at a concentration of 1000 cells/µl in PBS, 0.04% BSA for library preparation and sequencing.

### Single-cell RNA sequencing

Libraries were generated using the Chromium Next GEM Single Cell 3’ Reagent Kits v3.1 Dual Index (10X Genomics Inc., USA). Briefly, a droplet emulsion targeting 5,000 cells was generated in a microfluidic Next GEM Chip G, followed by barcoded cDNA generation inside the droplets. Purified and amplified cDNA was then subjected to library preparation and sequenced on a NovaSeq 6000 instrument (Illumina, USA) using a SP100 flow cell kit to a depth of 400M read-pairs per sample. Sequencing was performed as a service at the Department of Molecular Medicine (MOMA) of Aarhus University Hospital, Denmark (https://www.moma.dk/).

Single-cell reference annotation was carried out by the NGS service provider Omiics (https://omiics.com). In brief, barcode processing and single-cell 3′ gene quantification was done using the Cell Ranger Single-Cell Software Suite (v 3.1.0) using the GRCh38 reference genome. The Seurat R package (v 4.0.4) was used to quality filter cells and remove cells with above 20% of reads mapping to mitochondrial genes. Single cells were further filtered using the R package DoubletFinder (v.2.0.3) to remove the doublets. Seurat was used for data normalization, dimensionality reduction by use of the Uniform Manifold Approximation and Projection (UMAP) technique. Clustering was done with Seurat’s graph-based clustering approach using the FindClusters function. Annotation of clusters was assigned manually based on previously published data of single-cell sequencing using cortical differentiation (Camp et al., 2015; Eze et al., 2021) (Fleck et al., 2021; Ziffra et al., 2021).

### Statistical analysis

Statistical analysis was performed using GraphPad Prism 9. Data are given as the mean ± standard deviation (SD). Statistical significance of data was determined by one-way ANOVA followed by Dunnett’s multiple comparisons tests.

## RESULTS AND DISCUSSION

### VPS10P domain receptors show temporally distinct expression patterns during organoid development

To interrogate expression of VPS10P domain receptors during human brain development and in adult function, we modified published protocols for efficient and reproducible generation of cerebral organoids from human iPSC lines (see methods for details). Our protocol follows a time line of seeding of iPSCs, formation of embryoid bodies (EB), and subsequent maturation of the organoids for up to 100 days (Fig. 1B). Cerebral organoids were chosen based on databases suggesting expression of all VPS10P domain receptors in human brain cortex (https://www.proteinatlas.org).

During maturation, cerebral organoids showed the expected decrease in pluripotency markers, such as *NANOG* and *OCT4*, as well as a corresponding induction of genes in early telencephalic development, including *PAX6* and *SOX2*, and an increase in the forebrain marker *FOXG1* (Fig. 2A). From day 14 onwards, marker gene expression indicative of neural development, including pan-neuronal markers *TUBB* and *MAP2*, as well as neuroepithelial marker *S100β* was apparent. Expression of non-neuronal cell markers *GFAP* (astrocytes) and *OLIG2* (oligodendrocytes) was seen in mature organoids around 100 days of age (Fig. 2B). Markers of cortical specification appeared around day 21, including *TBR2* (cortical neurogenesis), *VGLUT2* (glutamatergic neurons), as well as *CTIP2* and *TBR1* (cortical neurons layers V and VI) (Fig. 2C). Induction of neurogenesis around day 14 (Fig. 3A), and subsequent formation of neuronal and non-neuronal cell types of the cerebral cortex was confirmed by immunohistology (Fig. 3B).

**Figure 2:**
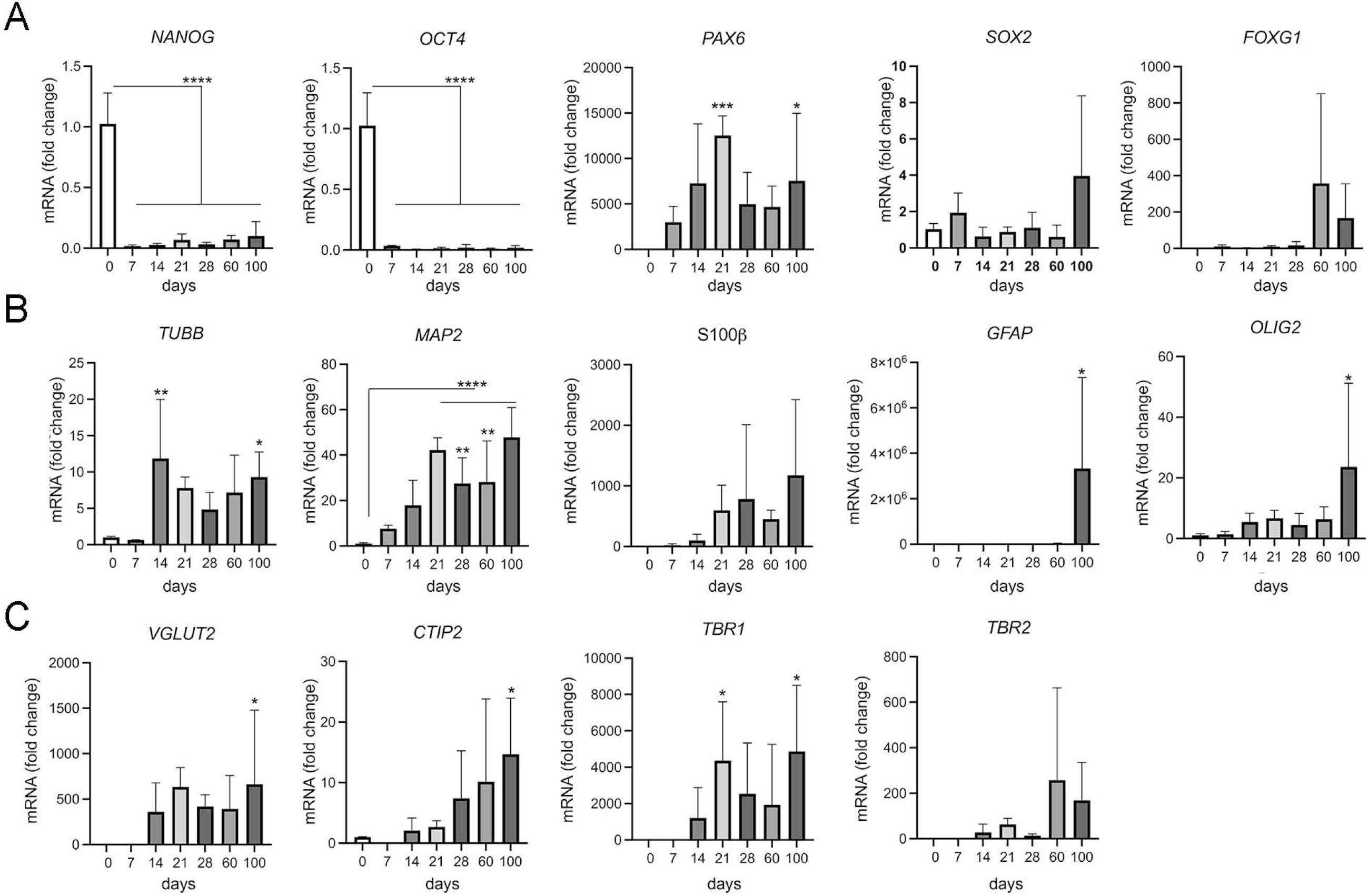
Marker gene transcription during organoid maturation. Transcript levels for the indicated marker genes were determined by quantitative (q) RT-PCR in cerebral organoids at various timepoints of differentiation. Data are given as changes relative to transcript levels in embryoid bodies (day 0 of differentiation set to 1). Statistical significance of data was determined by one-way ANOVA followed by Dunnett’s multiple comparisons tests. n = 3 - 5 organoids per condition; *, p<0.05; **, p<0.01; ***, p<0.001; ****, p<0.0001.

**Figure 3:**
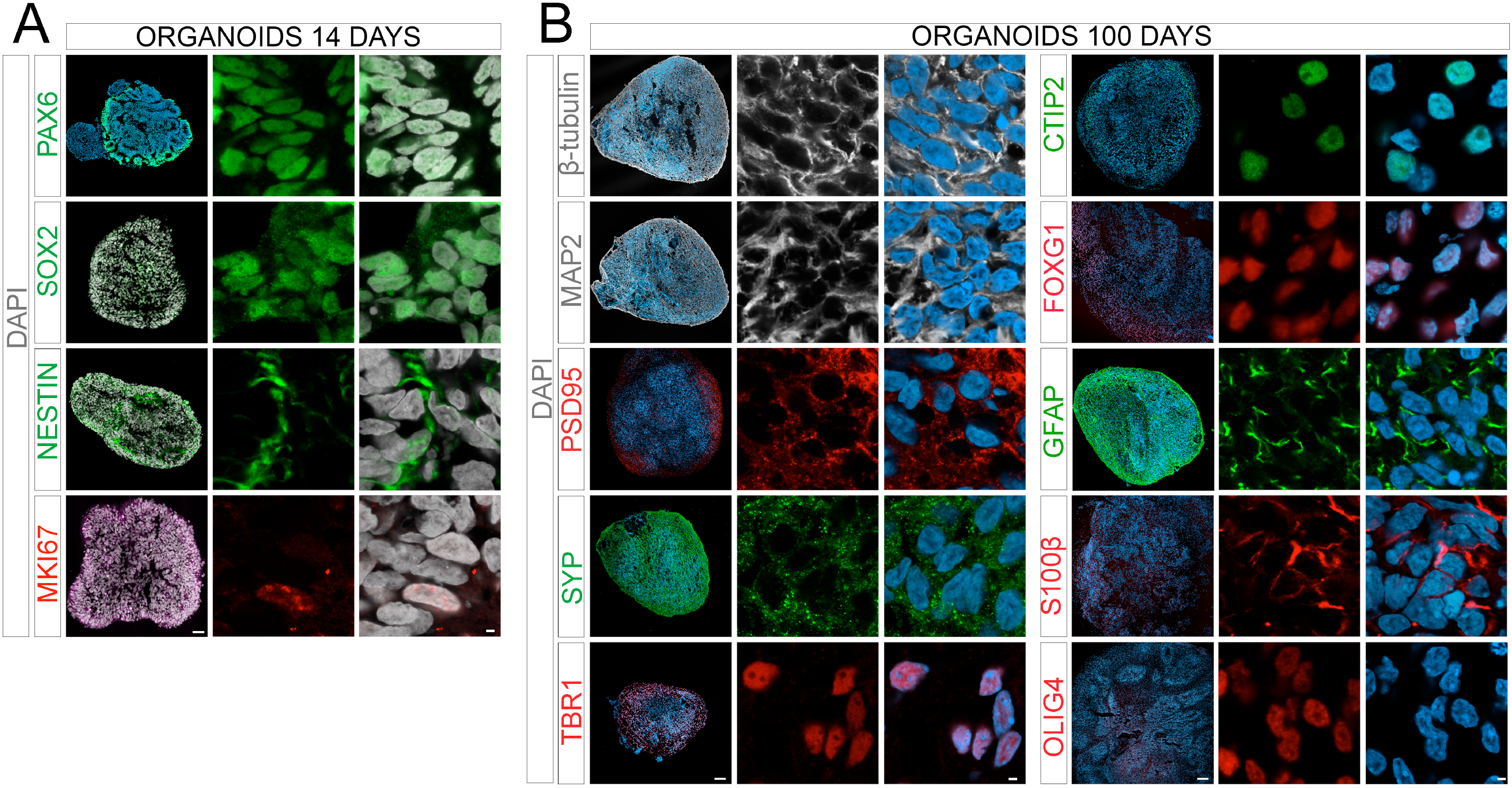
Marker protein expression in organoids at day 14 and 100. Immunohistological analysis of expression of the indicated marker proteins in cerebral organoids at 14 (**A**) or 100 days (**B**) of differentiation. Overview images in the left panels show merged channel configurations. Higher magnification images to the right show marker expression without (middle panels) or with nuclear DAPI stain (right panels). Scale bars: 2 µm or 100 µm (for overview images).

In line with expression of VPS10P domain receptors in murine (https://www.informatics.jax.org) and human brains (https://www.proteinatlas.org), human cerebral organoids showed induction of receptor gene expression. However, distinct differences were noted comparing receptor species (Fig. 4). Induction of gene transcription starting with early neurogenesis (day 14) was observed for *SORT1* (encoding sortilin) and *SORCS2*. A similar continuous increase in expression with organoid maturation was seen for *SORCS3*, although gene transcription was induced much later (from day 60) and remained overall low (Ct value 30.7 at day 100) as compared to *SORT1* (Ct value 23.1 at day 100) and *SORCS2* (Ct value 25.0 at day 100). By contrast, transcript levels for *SORL1* (encoding SORLA) were relatively high in EBs but continuously decreased with organoid formation (Ct value 27.1 at day 100). *SORCS1* transcripts showed a remarkable peak around day 21 and a subsequent decrease in levels in older organoids (Ct value 27.3 at day 100). Taken together, these data argued for distinct temporal functions of VPS10P domain receptors in the human brain with SORLA likely representing a marker of early progenitor fate, SORCS1 acting at a time of neuronal induction, while sortilin and SORCS2 representing markers of mature cerebral cell types. Low levels of *SORCS3* transcripts recapitulated the predicted insignificant expression of the receptor protein in human cortical tissue (https://www.proteinatlas.org).

**Figure 4:**
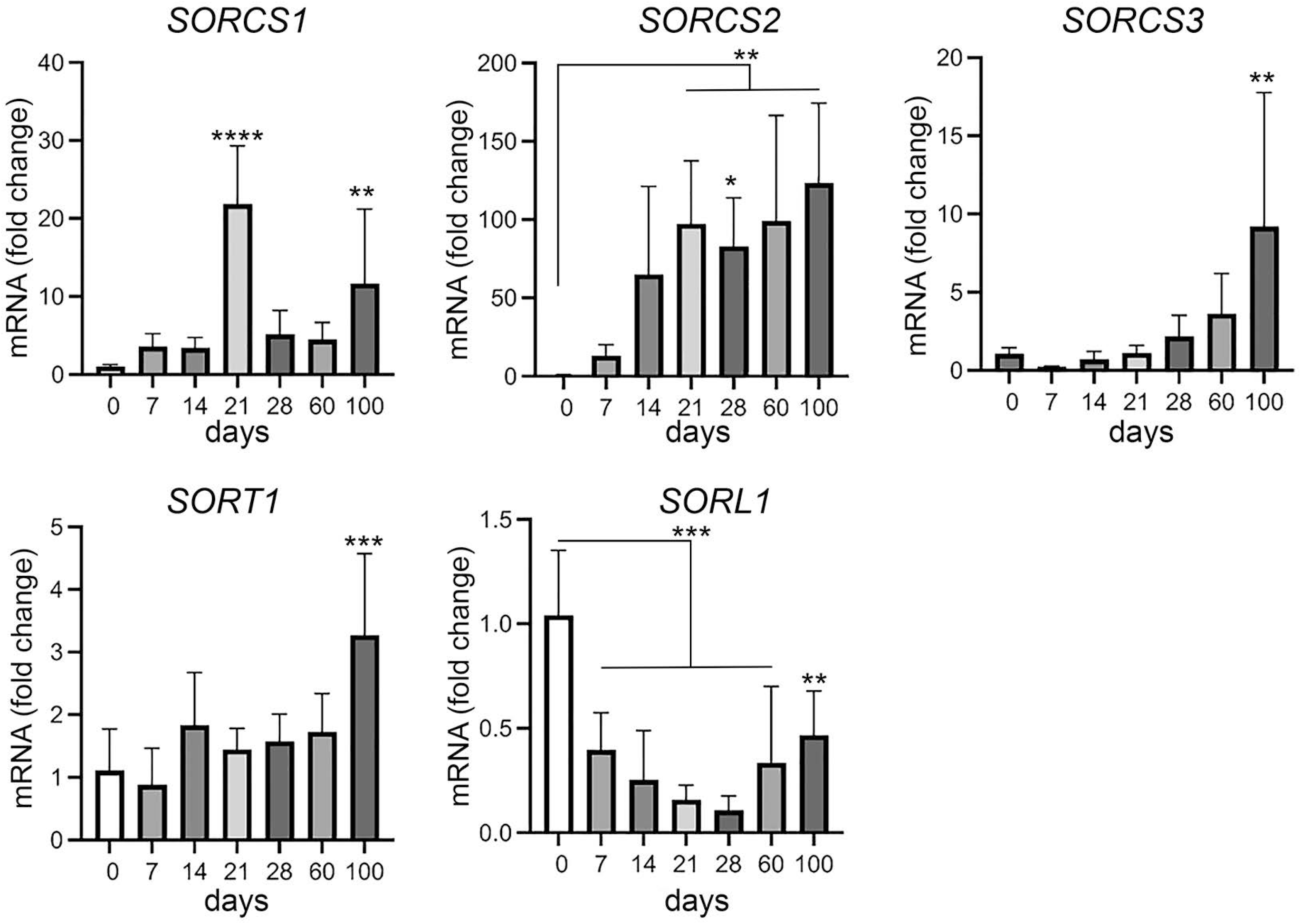
Transcript levels for VPS10P domain receptor genes during organoid maturation. Transcript levels for VPS10P domain receptor genes were determined by quantitative (q) RT-PCR in cerebral organoids at the indicated timepoints of differentiation. Data are given as changes relative to transcript levels in embryoid bodies (day 0 of differentiation set to 1). Statistical significance of data was determined by one-way ANOVA followed by Dunnett’s multiple comparisons tests. n = 4 - 13 organoids per condition; **, p<0.01; ***, p<0.001. ****, p<0.0001.

### Cerebral organoids express functionally distinct variants of VPS10P domain receptor species

While transcript levels constitute an accurate representation of receptor gene induction during cortical development, functional expression of VPS10P domain receptors is commonly regulated by post-transcriptional mechanisms, including glycosylation (Rovelet-Lecrux et al., 2021) or proteolytic processing into various receptor isoforms (Glerup et al., 2014). Therefore, we tested expression of VPS10P domain receptor isoforms in cerebral organoids by western blotting, documenting abundant expression of SORCS1 in early organoids at days 14 and 21 (Fig. 5A) and of all receptors, with the exception of SORCS3, at day 100 (Fig. 5B). In line with studies in mice and established cell lines (Hermey et al., 2003; Glerup et al., 2014), SORCS1 and SORCS2 were present as several receptor isoforms as seen from multiple immunoreactive bands in the 130-150 kDa molecular weight range (brackets in panel 5B). Also, detection of a 104 kDa protein band, representing the cleaved amino terminal domain of SORCS2 (asterisk in panel 5B), documented the presence of single- and two-chain variants of the receptor in human cerebral organoids. These two SORCS2 variants have been reported in murine cell types and shown to carry distinct functions in trophic support of dopaminergic innervation (single-chain form) and in induction of cell death in Schwann cells (two-chain form) (Glerup et al., 2014). In further support of data from rodent cell models, full-length SORCS1 in human cerebral organoids was undergoing proteolytic processing to release the soluble VPS10P domain of the receptor that acts as a diffusible factor in the circulation of mice to facilitate insulin receptor signaling (Kjolby et al., 2020). This fact was documented by immunoprecipitation of the SORCS1 ectodomain from the supernatant of organoids at days 14, 21, and 100 (Fig. 5C-D). Finally, soluble ectodomains of SORLA and sortilin, produced by membrane shedding (Hampe et al., 2000; Hermey et al., 2006), were also detectable in organoid cultures (Fig. 5E). Although the functional relevance of these receptor fragments remains poorly explained, they bear significance as surrogate biomarkers in blood or cerebrospinal fluid, mirroring levels of receptor expression in tissues (Ma et al., 2009; Carrasquillo et al., 2010). In conclusion, these findings identified the existence of biologically relevant receptor isoforms and processing products in human cerebral organoids, making them a useful model to study the roles of these receptor variants for human brain functions.

**Figure 5:**
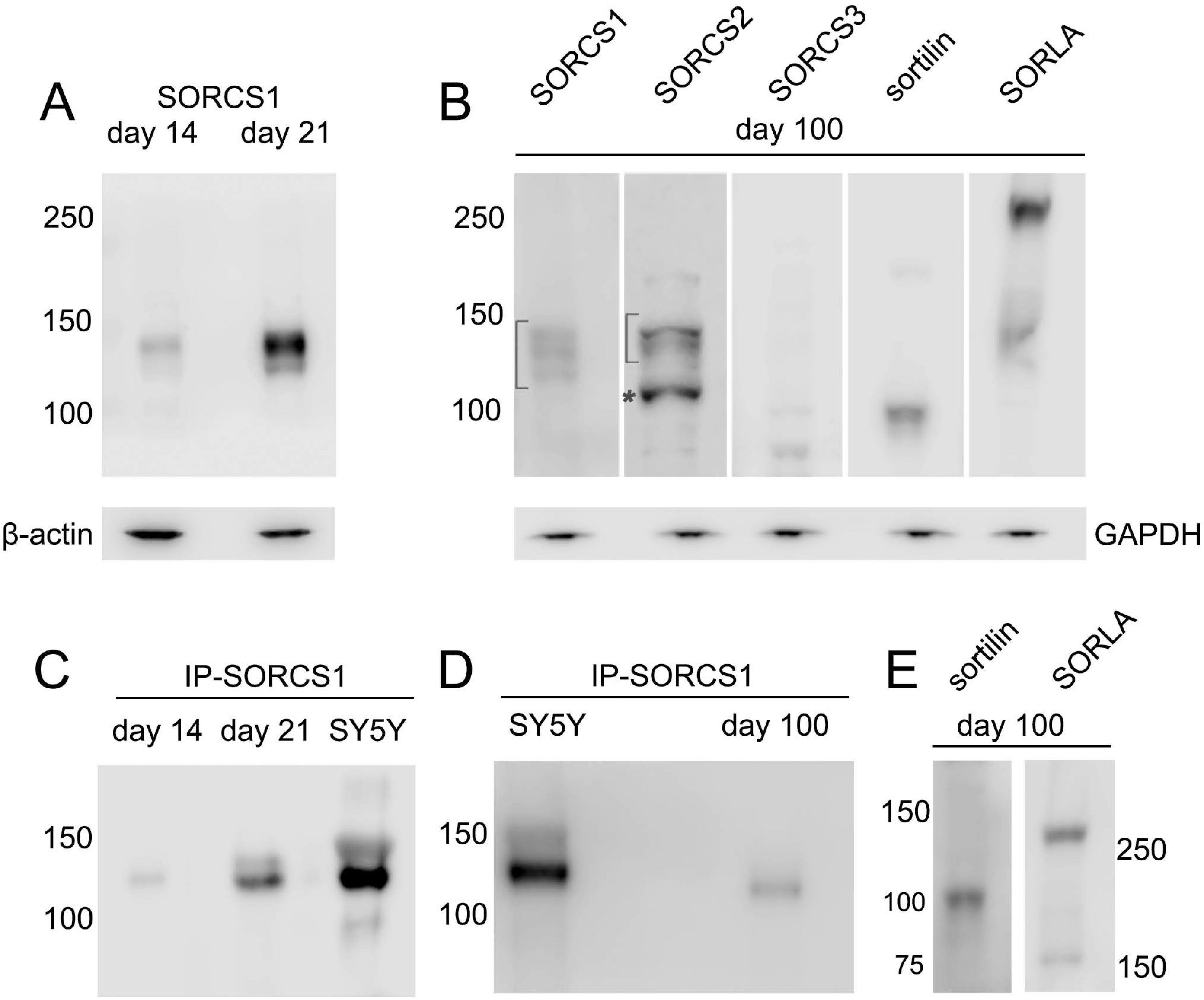
Expression of VPS10P domain receptors in cerebral organoids during maturation. (**A, B**) Expression of VPS10P domain receptors was tested in membrane protein preparations from cerebral organoids at 14 and 21 days (A) or 100 days (B) of differentiation using western blotting. In B, brackets indicated the range of migration of various isoforms of SORCS1 and SORCS2. The asterisk denotes the 104 kDa amino terminal fragment of the two-chain form of SORCS2. Detection of β-actin (**A**) and GAPDH (**B**) served as loading controls. (**C, D**) The soluble ectodomain of SORCS1 was recovered by immunoprecipitation (IP-SORCS1) from cytosolic extracts of cerebral organoids at day 14 and 21 (**C**) as well as day 100 (D) of differentiation. The SORCS1 ectodomain recovered by IP from the cell supernatant of the neuroblastoma cell line SY5Y is shown as positive control. (**E**) Immunodetection of shedded ectodomains of sortilin and SORLA in soluble extracts of day 100 organoids. For all panels, the migration of marker proteins of the indicated molecular weights in kDa are given.

### VSPS10P domain receptors show spatially distinct patterns of expression in neuronal and non-neuronal brain cell types

To explore the cell-type specific expression of VPS10P domain receptors in cerebral organoids, we performed single-cell RNA sequencing (scRNAseq) in mature organoids at day 100. To do so, we developed novel protocols to generate single-cell suspensions with reproducible cell viability of >80%, a technical obstacle that complicates the analyses of mature organoids (see methods for details). Analysis of transcript data by unsupervised clustering and visualized using Uniform Manifold Approximation and Projection for dimension reduction (UMAP) plot identified 18 distinct cell clusters in 100 days old cerebral organoids, including neuronal progenitors (e.g., clusters 2 and 13) as well as immature (e.g., clusters 3 and 5) and mature (e.g., clusters 0, 14, 15) neuronal cell types. Radial glia (cluster 12), mesenchymal cells (cluster 16), and astrocytes (cluster 8) were among the non-neuronal cell types identified (Fig. 6A and Suppl. figure S1). Amongst the VPS10P domain receptors, *SORT1* showed the most wide-spread robust expression. Transcripts were observed in neuronal progenitors and immature and mature neuronal cell types, but also in radial glia and astrocytes (Fig. 6B). *SORL1* transcripts were also wide-spread but less abundant than those for *SORT1* (Fig. 6B). *SORCS1* transcript levels were low in aged organoids and restricted to neuronal progenitors and astrocytes (Fig. 6C). *SORCS2* transcripts were seen in mature neuronal and non-neuronal clusters, while *SORCS3* transcripts were rarely detectable (Fig. 6C). These data substantiated conclusions from qRT-PCR, associating *SORL1* with progenitor cell functions, *SORCS1* with early neurogenesis, and *SORCS2* and *SORT1* with more mature brain cell types. Of note, the inherent lack of microglia in cerebral organoids precluded us from substantiating the expression of VPS10P domain receptors, notably *SORL1*, in this cell type. However, we have reported *SORL1* expression in iPSC-derived human microglia using specialized differentiation protocols before (Kaminska et al., 2023).

**Figure 6:**
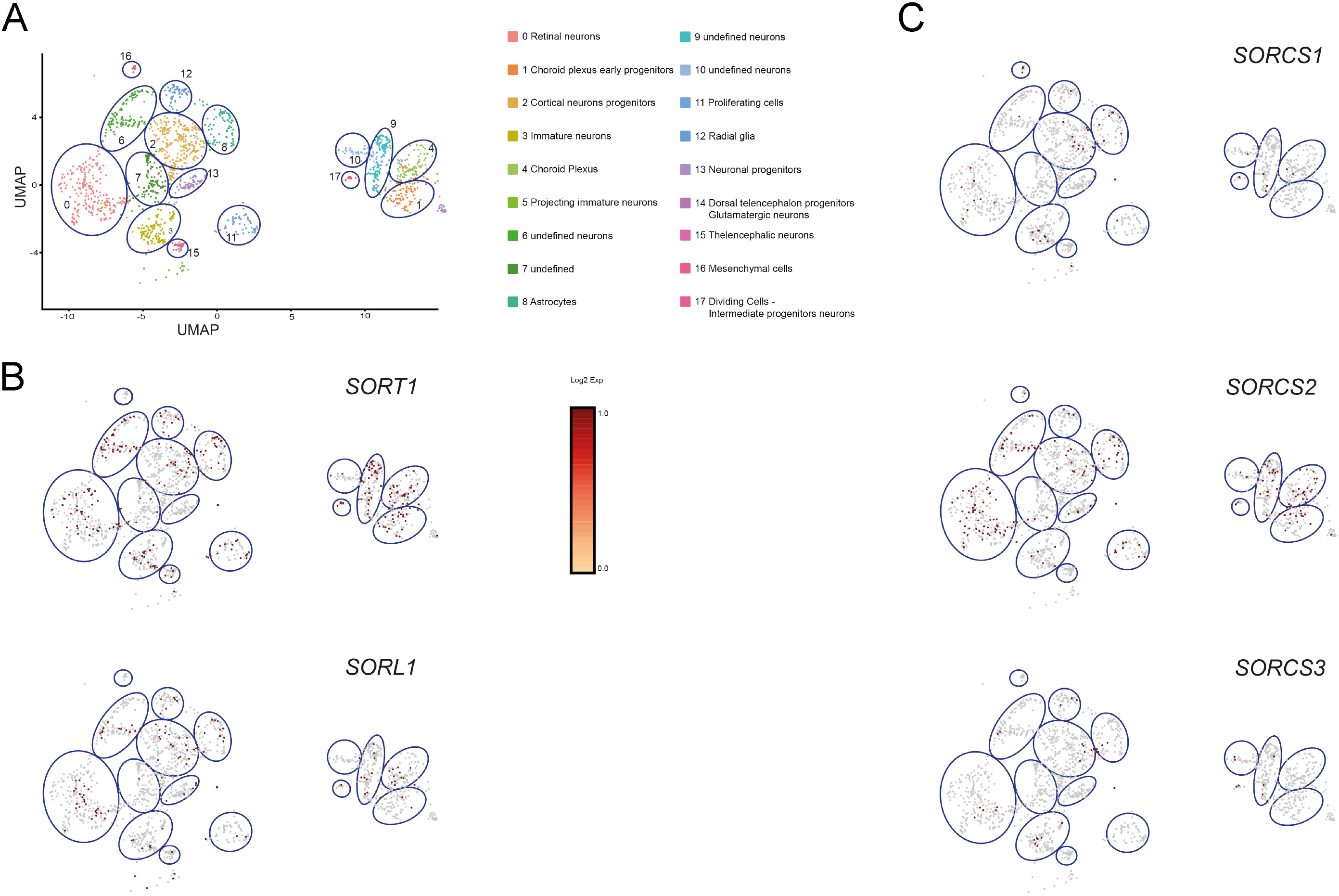
Brain cell-type specific expression of VPS10P domain receptors identified by single-cell RNAseq. (**A**) Cells from100 days old cerebral organoids were analyzed by unsupervised clustering and visualized using Uniform Manifold Approximation and Projection for dimension reduction (UMAP) plot. A total of 18 distinct cell clusters were identified in the data set. (**B-C**) UMAP plots localizing VPS10P domain receptor transcripts to the indicated clusters.

Cell-type specific expression of VPS10P domain receptors in cerebral organoids was corroborated by immunohistochemistry. At day 14, expression of SORCS1 was localized to proliferating neuronal progenitors characterized by expression of NESTIN, PAX6 and SOX2, as well as proliferation marker MKI67 (Fig. 7A). At day 100 of maturation, robust levels of SORCS1, SORCS2, and sortilin were detected in cortical neurons expressing MAP2, TBR1, CTIP2, and VGLUT (Fig. 7B). Despite a substantial drop in transcript levels in organoids as compared to EBs, SORLA protein was easily detectable in cortical neurons at day 100 of maturation (Fig. 7B). Immunohistochemistry recapitulated detection of the receptor by western blotting (Fig. 5B). This finding is noteworthy as prior scRNAseq analyses suggested that expression of this AD risk factor in the human brain may be restricted to microglia (Gosselin et al., 2017; Hansen et al., 2018). However, based on data in this study, and in line with immunohistological analyses of human brain specimens (Scherzer et al., 2004; Offe et al., 2006), substantial levels of SORLA can clearly be seen in human neurons in cerebral organoids. In addition to neuronal cell types, SORLA, sortilin, and SORCS1 were also expressed in GFAP-positive astrocytes in 100 days old organoids (Fig. 7B). Lower astrocytic expression was seen for SORCS2 (Fig. 7B), corroborating findings from mouse models that expression of this receptor in astrocytes is only induced during stress response, as in stroke (Malik et al., 2020).

**Figure 7:**
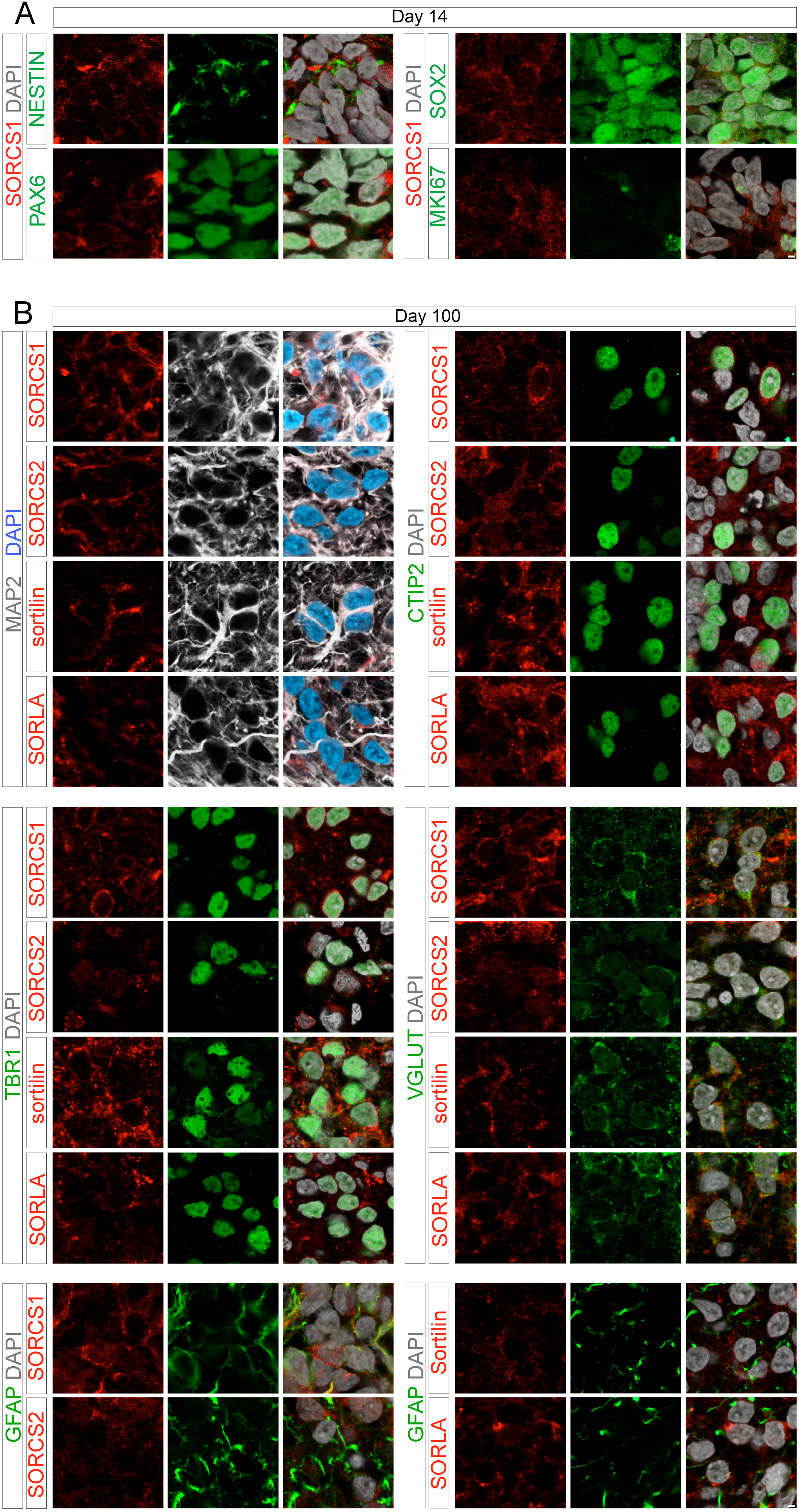
Cell-type specific expression of VPS10P domain receptors in 14 and 100 days old organoids. Immunohistological detection of the indicated VPS10P domain receptors in 14 day (**A**) or 100 day (**B**) old cerebral organoids. Cell-type specific expression of the receptors (red) was tested by colocalization with the indicated cell markers (green, grey). Single (left, middle) as well as merged channel configurations with DAPI counterstain (right) are shown. Scale bars: 2 µm.

### Subcellular localization supports distinct roles for VSPS10P domain receptors in intraneuronal protein transport

VPS10P domain receptors are mainly recognized for their role in sorting target proteins between cell surface and intracellular compartments, defining neuronal activities relevant for endocytosis, secretion, or signal reception. For example, SORLA has been identified as a sorting receptor for the amyloid precursor protein (APP), moving internalized APP from endosomes to the Golgi to prevent endosomal breakdown into amyloid-β peptides (Andersen et al., 2005; Burgert et al., 2013; Dumanis et al., 2015). Sortilin acts as an endocytic receptor for apolipoprotein E4 and progranulin, etiologic agents in AD and FTD, respectively (Hu et al., 2010; Asaro et al., 2020). SORCS1, -2, and -3 sort the neurotrophin receptor tropomyosin receptor kinase (Trk) B to synaptosomal membranes, controlling neurotrophic signal reception in target cells (Glerup et al., 2016; Subkhangulova et al., 2018). To explore distinct subcellular localization patterns in human cerebral neurons, we co-immunostained VPS10P domain receptors with markers of various intracellular compartments in 100 days old organoids (Fig. 8). Abundant expression of all tested receptors was observed in cortical neurons, yet differences in subcellular localization were apparent. While SORLA mainly localized to the perinuclear region and intracellular vesicles in the soma, strong SORCS1 immunoreactivity was seen in the soma but also in neuronal projections (arrowheads in Fig. 8). Like SORLA, sortilin mainly localized to the perinuclear region, however some immunofluorescence signal was also present in neuronal projections (arrowheads in Fig. 8). SORCS2 showed a distinct punctate pattern both in somatic vesicles but also in neuronal projections (Fig. 8).

**Figure 8:**
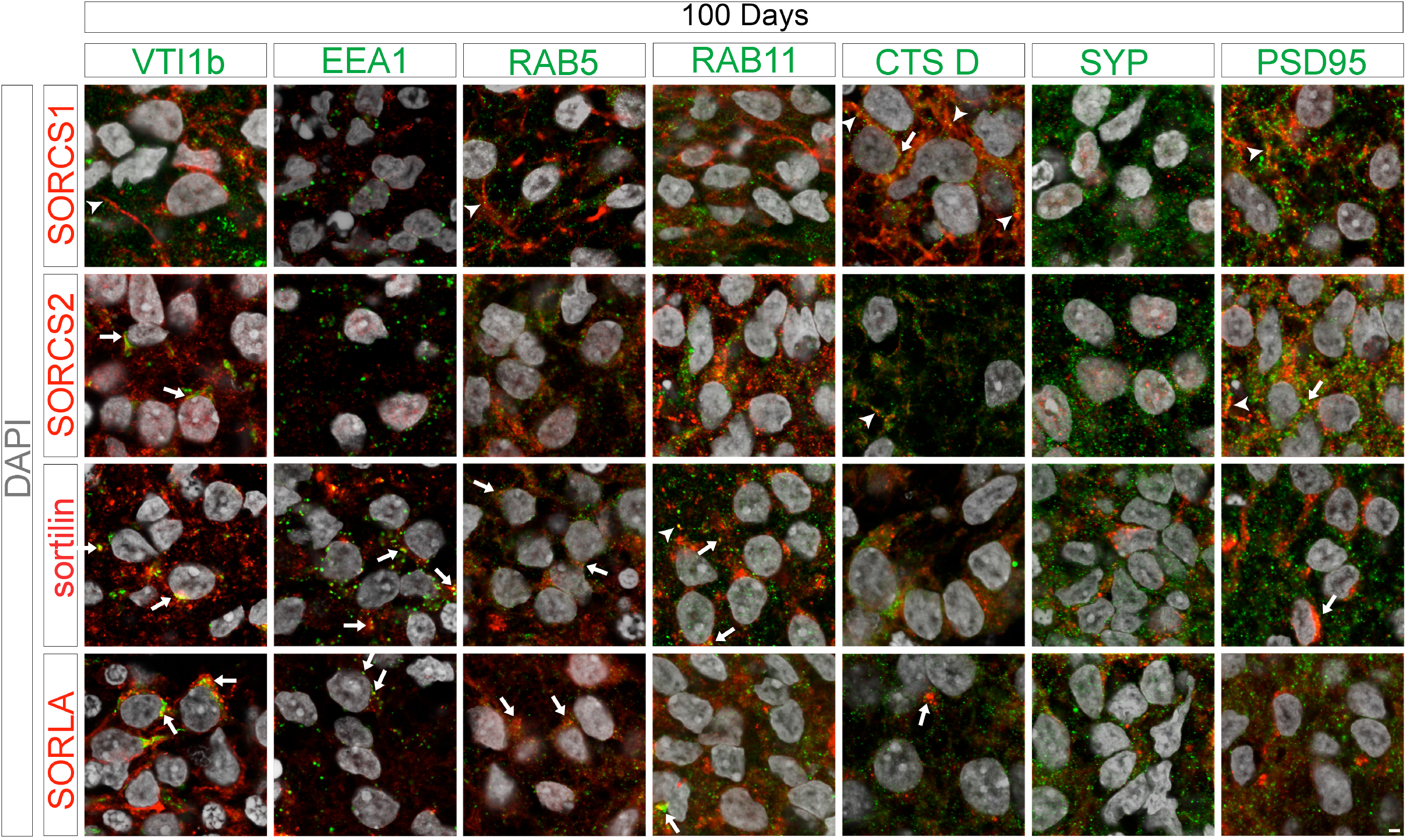
Subcellular localization of VPS10P domain receptors in 100 days old organoids. Colocalization of SORCS1, SORCS2, sortilin, and SORLA (red) with the indicated organelle markers (green) was tested using immunohistochemistry. Merged channel configurations including nuclear counterstain with DAPI (grey) are shown. Arrowheads mark the localization of individual receptors to neuronal projections in exemplary images. Arrows highlight co-localization of the VPS10P domain receptors with the given markers. Scale bars: 2 µm.

Co-staining with markers refined the localization of VPS10P domain receptors to distinct subcellular compartments in human cortical neurons (Fig. 8). In line with abundant localization to the perinuclear region, SORLA and sortilin co-localized with the Golgi marker VTI1b. Receptor-containing somatic vesicles encompassed early (EEA1, Rab5) and recycling (Rab11) endosomes. SORLA, but not sortilin, also localized to lysosomes, stained for cathepsin D. Overall, these patterns support established roles for both receptors in endocytosis as well as Golgi to endo-lysosomal sorting. SORCS1 and SORCS2 mainly localize to the Golgi and to Rab5-positive endosomes, but also to lysosomes (Cathepsin D) in the soma and along neuronal projections. Although speculative at present, this path would be consistent with roles for these receptors in retrograde sorting of signaling endosomes containing activated Trk receptors and possible involvement in lysosomal protein degradation in axons (Yamashita and Kuruvilla, 2016; Ferguson, 2018). Given their roles in synaptic plasticity, e.g., by controlling brain-derived neurotrophic factor signaling through TrkB (Vaegter et al., 2011; Glerup et al., 2016; Subkhangulova et al., 2018), sortilin as well as SORCS1 and -2 also colocalized to the post-synaptic density marked by PSD95. By contrast, receptor immunoreactivity was absent from the pre-synaptic compartment identified by synaptophysin (Fig.8). No obvious localization of SORLA to pre- or post-synaptic compartments was observed.

In conclusion, our studies uncovered distinct temporal, spatial, and subcellular expression patterns for VPS10P domain receptors in human cerebral organoids. The observed patterns argue for divergent roles for these receptors in development and adult brain function, and for distinct modes of action in sorting paths in soma and somatodentritic compartments of human neurons. Importantly, our data suggest distinct contributions of *SORCS1*, but also *SORL1*, to neuronal progenitor fate and function during early neurogenesis. Such activities may explain association of *SORCS1* with neurodevelopmental disorders, such ADHD (Lionel et al., 2011). Based on our data, cerebral organoids derived from iPSC lines of patients and control subjects represent ideal model systems to further elucidate the pathophysiological mechanisms underlying such receptor (dys)functions in the human brain.

## Acknowledgment

We are indebted to Kristin Kampf and Anne Højland for expert technical assistance, to Dr. Silke Frahm-Barske (MDC) for sharing protocols, and to Prof. Anders Nykjaer (Aarhus University) for critical reading of the manuscript. Studies were funded in part by grants from the Novo Nordisk Foundation (NNF18OC0033928) to TEW.

## Data availability

All data are available on reasonable request from the authors. The scRNAseq data are available from the GEO database.

## Authors’ relationships and activities

The authors declare no competing interest related to this manuscript.

## Contribution statement

FF conceptualized the study, performed experiments, and analyzed data. HHBA performed experiments and analyzed data. TEW conceptualized the study and interpreted data. FF and TEW wrote the manuscript.

