## Supplementary material for "Spatially and temporally distinct patterns of expression for VPS10P domain receptors in human cerebral organoids"

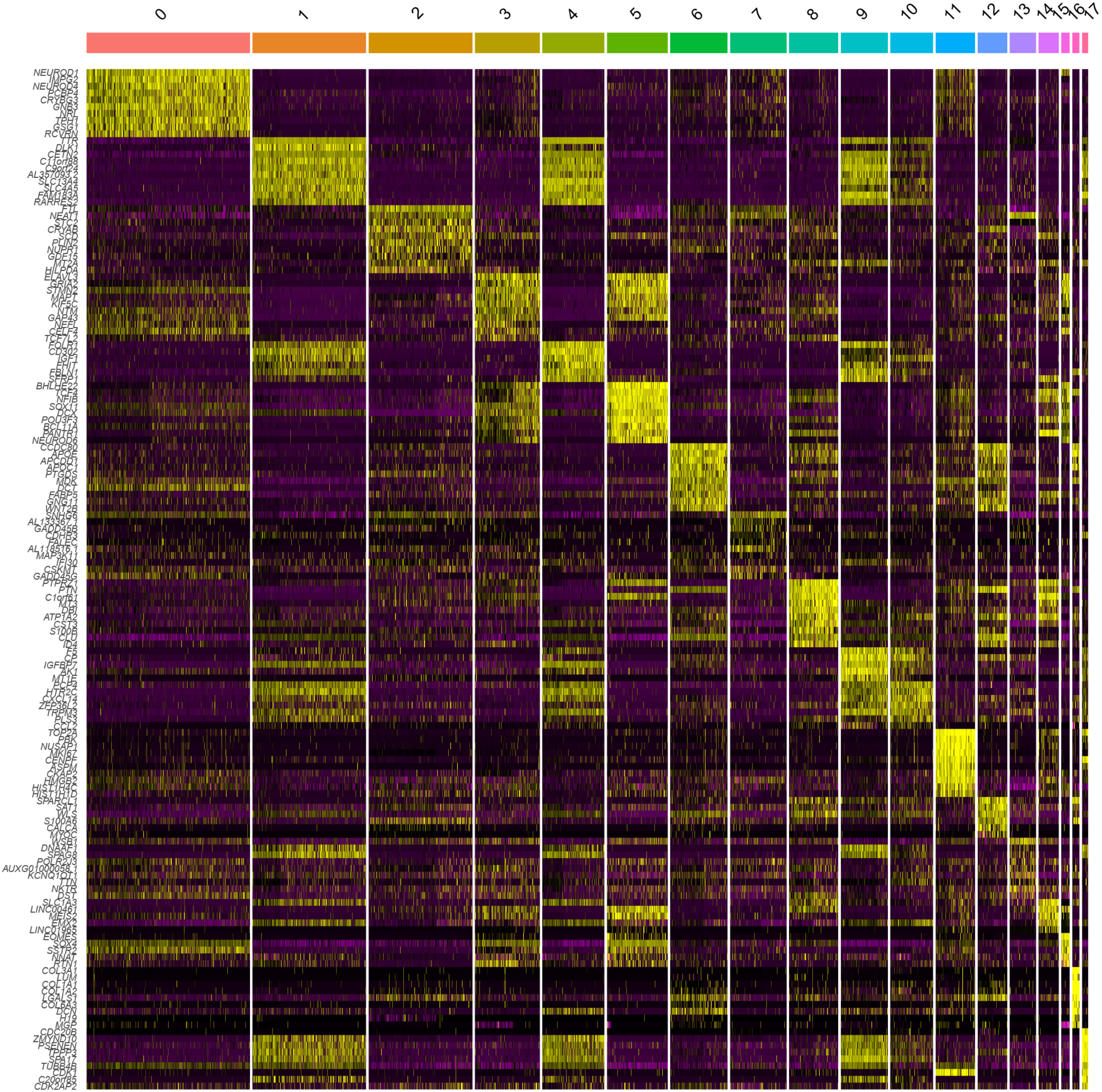
NEUROD1
IMPG2
NEUROD4
PCBP4
CRYBG3
GNB3
GSG1
RCVRN
TTR
DLK1
C11orf88
C9orf24
AL357093.2
SLC13A4
RARRES2
FTL
NEAT1
STC2
CRYAB
PLIN2
NUPR1
GDF15
MT2A
HILPDA
STMN2
MAPT
KIF5C
NTM
GAP43
NEFL
CELF4
TCF7L2
FOLR1
CD302
IGF1
FBLN1
SFRP1
BHLHE22
TCF4
NFIB
SOX11
POU3F3
BCL11A
PANTR1
NEUROD6
APOE
APCDD1
APOC1
PTGDS
MDK
FABP5
GNG11
WNT2B
SNHG6
GADD45B
AL118516.1
MAP3K11
IFI30
GADD45G
PTPRZ1
PTN
C1orf61
MT3
ATP1A2
CST3
S100B
CLU
ID4
IGFBP7
AK1
MT1E
HTR2C
ZFP36L2
TRPM3
PLS3
TOP2A
NUSAP1
MKI67
CENPF
ASPM
HMGB2
HIST1H4C
HIST1H1D
SPARCL1
S100A6
CALCA
MYOC
WSB1
DNAAF1
AUXG01000058.1
KCNQ1OT1
TTN
NKTR
SLC1A3
LINC00461
MEIS2
EMX2
LINC01965
EOMES
SSTR2
NNAT
RTN1
COL3A1
LUM
COL1A1
COL1A2
LGALS1
COL6A3
DCN
MGP
CDC20B
ZMYND10
PSENEN
TPPP3
SPA17
TUBB4B
C20orf85
CDK2AP2

**SUPPLEMENTARY TABLES**

### Supplememtary table 1: Taqman probes used for qRT-PCR

| **Primer** | **Manufacturer** | **Catalog number** | **Species** |
|---|---|---|---|
| *CTIP2 (BCL11B)* | Thermo Scientifics | Hs01102259_m1 | Human |
| *FOXG-1* | Thermo Scientifics | Hs01850784_s1 | Human |
| *GAPDH* | Thermo Scientifics | Hs99999905_m1 | Human |
| *GFAP* | Thermo Scientifics | Hs00909233_m1 | Human |
| *MAP2* | Thermo Scientifics | Hs00258900_m1 | Human |
| *NANOG* | Thermo Scientifics | Hs02387400_g1 | Human |
| *OLIG2* | Thermo Scientifics | Hs00300164_s1 | Human |
| *OTX2* | Thermo Scientifics | Hs00222238_m1 | Human |
| *PAX6* | Thermo Scientifics | Hs00242217_m1 | Human |
| *POU5F1 (OCT4)* | Thermo Scientifics | Hs00999632_g1 | Human |
| *S100B* | Thermo Scientifics | Hs00902901_m1 | Human |
| *SorCS1* | Thermo Scientifics | Hs00364666_m1 | Human |
| *SorCS2* | Thermo Scientifics | Hs01019137_m1 | Human |
| *SorCS3* | Thermo Scientifics | Hs01039447_m1 | Human |
| *SORL* | Thermo Scientifics | Hs00983772_m1 | Human |
| *SORT1* | Thermo Scientifics | Hs00361760_m1 | Human |
| *SOX2* | Thermo Scientifics | Hs01053049_s1 | Human |
| *TBR-1* | Thermo Scientifics | Hs00232429_m1 | Human |
| *TBR-2 (EOMES)* | Thermo Scientifics | Hs00172872_m1 | Human |
| *TUBB3* | Thermo Scientifics | Hs00964963_g1 | Human |
| *VGLUT2 (SLC17A6)* | Thermo Scientifics | Hs00220439_m1 | Human |

### Supplementary table 2: Primary antibodies used in this study

| **Antibody** | **Manufacturer** | **Catalog number** | **Host** | **Dilution** |
|---|---|---|---|---|
| β–actin | Abcam | AB8227 | Rabbit | 1:1000 |
| Cathepsin D | R&D System | MAB10141 | Mouse | 1:500 |
| CTIP2 | Abcam | AB18465 | Rat | 1:500 |
| NESTIN | ABCAM | AB105389 | Rabbit | 1:200 |
| EEA1 | BD Biosciences | 610457 | Mouse | 1:200 |
| FOXG1 | Abcam | AB227888 | Rabbit | 1:100 |
| GAPDH | Genetex | GTX627408-01 | Mouse | 1:2000 |
| GFAP | Abcam | AB53554 | Goat | 1:200 |
| Ki67/MKI67 | R&D System | AF7617 | Sheep | 1:40 |
| MAP2 | Synaptic System | 188004 | Guinea pig | 1:1000 |
| OLIG4-PE | Melteny | 130-109-199 | Mouse | 1:10 |
| PAX6 | Biolegend | 901301 | Rabbit | 1:400 |

| | | | | |
|---|---|---|---|---|
| PSD 95 | Abcam | AB2723 | Mouse | 1:200 |
| RAB5A | Synaptic System | 108011 | Mouse | 1:500 |
| RAB11 | Cell Signaling | 5589 | Rabbit | 1:50 |
| SORCS1 | Sigma | HBA011948 | Rabbit | 1:250<br>(4µl for IP) |
| SORCS1 | R&D System | AF3457 | Goat | 1:70 (IF)<br>1:1000 (WB) |
| SORCS2 | DAKO F7100 | | Rabbit | 1:200 (IF)<br>1:1000 (WB) |
| SORCS2 | R&D System | AF4238 | Sheep | 1:40 (IF) |
| SORCS3 | R&D System | MAB30671 | Mouse | 1:1000 (WB) |
| SORLA | BD Bioscience | 611861 | Mouse | 1:1000 (WB)<br>1:1000 (IF) |
| SORLA | in house | Mouse | Goat | 1:40 |
| sortilin | BD bioscience | BD612101 | Mouse | 1:1000 (WB) |
| sortilin | R&D | AF3154 | Goat | 1:200 |
| SOX2 | R&D System | MAB2018 | Mouse | 1:500 |
| Synaptophysin | ENZO LIFE SCI | ADI-905-782 | Rabbit | 1:1000 |
| S100β | Dako | GA504 | Rabbit | 1:200 |
| TBR1 | Abcam | AB31940 | Rabbit | 1:200 |
| β-tubulin | Merk | MAB1637 | Mouse | 1:1000 |
| VGLUT | Synaptic System | 135403 | Rabbit | 1:500 |
| VTi1b | BD Biosciences | 611405 | Mouse | 1:150 |
| DAPI | Invitrogen | D1306 | | 1:1000 |

**Supplementary table 3: Secondary antibodies used in this study**

| **Antibody** | **Manufacturer** | **Catalog number** | **Host** | **Dilution** |
|---|---|---|---|---|
| anti-mouse Alexa Fluor 488 | Invitrogen | A21202 | Donkey | 1:1000 |
| anti-mouse Alexa Fluor 568 | Invitrogen | A10037 | Donkey | 1:1000 |
| anti-mouse Alexa Fluor 647 | Invitrogen | A31571 | Donkey | 1:1000 |
| anti-goat Alexa Fluor 568 | Invitrogen | A11057 | Donkey | 1:1000 |
| anti-goat Alexa Fluor 568 | Invitrogen | A21447 | Donkey | 1:1000 |
| anti-rabbit Alexa Fluor 488 | Invitrogen | A21206 | Donkey | 1:1000 |
| anti-rabbit Alexa Fluor 568 | Invitrogen | A10042 | Donkey | 1:1000 |
| anti-rabbit Alexa Fluor 647 | Invitrogen | A31573 | Donkey | 1:1000 |
| anti-guinea Pig Alexa Fluor 647 | Jackson Immuno | 706-605-158 | Donkey | 1:200 |

### SUPPLEMENTARY METHODS

#### Expression analyses

Transcript levels were determined by quantitative (q) RT-PCR. To do so, total RNA was extracted from organoids using RNeasy Mini Kit (70104, Qiagen) and reverse transcribed using High-Capacity RNA to cDNA kit (4387406, Thermo Fisher Scientific). Quantitative PCR reaction was performed using 5 ng of cDNA and the Taqman Fast Advanced Master Mix solution (4444557, Thermo Fisher Scientific). The samples were loaded on a MicroAmp EnduraPlate (4483285, Life and technologies) and run on the QuantStudio 7 Flex Real-Time PCR System instrument (Thermo Fisher Scientific). The probes used are given in Suppl. table 1. Expression values were normalized to *GAPDH* as internal control. Protein levels were determined by standard western blot analyses using primary and secondary antibodies given in Suppl. tables 2 and 3.

#### Immunohistochemical analyses

Immunohistochemistry was performed on organoids fixed in 4% paraformaldehyde for 20 min, infiltrated in 20% sucrose over night at 4°C, and embedded in OCT compound (Tissue-Tek, Sakura) for standard 10 µm cryo-sectioning. For immunodetection, cryosections were washed with PBS for 10 min, blocked in 5% donkey serum (D9663, Sigma) in PBT (PBS with 0.25% Triton X-100) for 1 hour at room temperature, followed by incubation with primary antibody in 2,5% donkey serum in PBT at 4°C overnight. Antibodies are listed in Suppl. table 2.  Next, tissue sections were washed 3x with PBS for 10 minutes, blocked with 1% donkey serum in PBST for 10 minutes, before treatment with secondary antibodies (1% donkey serum in PBT), for 1 hour at 4°C. The secondary antibodies used are listed in the Suppl. table 3.  Finally, coverslips were washed in PBS and mounted on slides using Fluorescence Mounting Medium (S3023, Dako). Images were captured on a confocal microscope (Zeiss LSM800) using 10X and 63X lens.

### SUPPLEMENTARY FIGURES

**Supplementary figure 1:** Heatmap showing the top ten most unique identifiers per cluster in the single-cell RNAseq data set. Genes were identified using the Seurat function "FindAllMarkers" and visualized using the Seurat function "DoHeatmap".
